# Force generation of KIF1C is impaired by pathogenic mutations

**DOI:** 10.1101/2021.06.30.450611

**Authors:** Nida Siddiqui, Daniel Roth, Algirdas Toleikis, Alexander J. Zwetsloot, Robert A. Cross, Anne Straube

## Abstract

Intracellular transport is essential for neuronal function and survival. The fastest neuronal transporter is the kinesin-3 KIF1C. Mutations in KIF1C cause hereditary spastic paraplegia and cerebellar dysfunction in human patients. However, neither the force generation of the KIF1C motor protein, nor the biophysical and functional properties of pathogenic mutant proteins have been studied thus far.

Here we show that full length KIF1C is a processive motor that can generate forces up to 5.7 pN. We find that KIF1C single molecule processivity relies on its ability to slip and re-engage under load and that its slightly reduced stall force compared to kinesin-1 relates to its enhanced probability to backslip. Two pathogenic mutations P176L and R169W that cause hereditary spastic paraplegia in humans maintain fast, processive single molecule motility in vitro, but with decreased run length and slightly increased unloaded velocity compared to the wildtype motor. Under load in an optical trap, force generation by these mutants is severely reduced. In cells, the same mutants are impaired in producing sufficient force to efficiently relocate organelles.

Our results establish a baseline for the single molecule mechanics of Kif1C and explain how pathogenic mutations at the microtubule-binding interface of KIF1C impair the cellular function of these long-distance transporters and result in neuronal disease.

## Introduction

Intracellular transport along microtubules allows the distribution of membrane organelles, and shuttling building blocks and signalling molecules between the nucleus and the cell periphery. The transport is mediated my molecular motor proteins, dynein and kinesins that hydrolyse ATP to step unidirectionally along microtubules. While dynein is the major minus end-directed transporter, more than 20 mammalian kinesins mediate plus-end directed transport (Paschal and Vallee, 1987, Lipka et al., 2016) These translocating kinesins share a conserved N-terminal motor domain of about 350 amino acid length that binds to microtubules and ATP. ATP hydrolysis and microtubule binding affinity are tightly coupled so that kinesins hydrolyse one ATP for each 8nm step along the microtubule (Svoboda et al., 1993, Coy et al., 1999). The stalk and tail of translocating kinesins are diverse and mediate cargo specificity and the regulate motor activity through inhibitory intramolecular interactions (Friedman and Vale, 1999, Kaan et al., 2011, Yamada et al., 2007, Siddiqui et al., 2019). Perturbed intracellular transport results in neurodegenerative disorders. For example, mutations in three kinesins have been implicated in causing hereditary spastic paraplegia (HSP) and spastic ataxias: KIF5A (SPG10), KIF1A (SPG30) and KIF1C (SPG58) (Reid et al., 2002, Bouslam et al., 2007, Klebe et al., 2012, Gabrych et al., 2019). These disorders present with progressive lower limb spasticity and gait difficulties due to the degeneration of motor neurons in the corticospinal tract (Schule et al., 2016, Hedera et al., 2005, Novarino et al., 2014). Complicated forms of HSP, such as those caused by KIF1C loss of function mutations, present with additional neurological defects such as cerebellar dysfunction, dystonic tremor and demyelinating neuropathy (Marchionni et al., 2019, Dor et al., 2014, Caballero Oteyza et al., 2014, Yucel-Yilmaz et al., 2018). The disease-causing mutations are either located within the KIF1C motor domain and predicted to modify either microtubule binding or nucleotide hydrolysis or result in truncations or instability of the protein through missense mutations (Gabrych et al., 2019, Dor et al., 2014, Caballero Oteyza et al., 2014, Yucel-Yilmaz et al., 2018). Heterozygote mutations usually present with a subclinical phenotype, thus homozygosity or a combination of two different mutations is required for full presentation of SPG58. For example, carriers of KIF1C_G102A_, which modifies the highly conserved nucleotide binding p-loop, and KIF1C_P176L_, which is predicted to modify microtubule binding, present with early onset disease (Caballero Oteyza et al., 2014). Expression in COS-7 African green monkey cells revealed that KIF1C_G102A_ mislocalised to the perinuclear region as opposed to the normal accumulation at the cell periphery, supporting the idea that the mutation perturbs motility. KIF1C_P176L_ localized correctly, but while co-expression of wildtype KIF1C could rescue the perinuclear mislocalization of KIF1C_G102A_, KIF1C_P176L_ could not (Caballero Oteyza et al., 2014). This led to the hypothesis that P176L results in a weakened KIF1C motor. A mutation in a nearby residue, R169W, was identified as a homozygotic mutation in patients with childhood onset cerebellar ataxia (Dor et al., 2014) for which any functional studies are lacking to date.

The kinesin-3 KIF1C is a ubiquitously expressed transporter responsible for Golgi to ER trafficking and maintenance of Golgi structure, the delivery of integrins, which is important for cell polarity and adhesion, and the transport of secretory vesicles in neurons (Dorner et al., 1998, Lee et al., 2015, Theisen et al., 2012, Schlager et al., 2010). KIF1C is highly processive, can equally efficiently transport cargo into axons and dendrites and is the most efficient plus end-directed translocator (Lipka et al., 2016, Siddiqui et al., 2019). In contrast to other kinesin-3s, KIF1C is a stable dimer whose autoinhibition is controlled by an intramolecular interaction of its stalk domain with the microtubule binding interface of the motor domain. Upon binding of pseudo-phosphatase PTPN21 or cargo adapter Hook3 to the stalk, the motor domain is released and can engage with microtubules (Siddiqui et al., 2019). Hook3 can simultaneously interact with KIF1C and dynein to form opposite polarity motor complexes (Kendrick et al., 2019), which might explain why depletion of KIF1C causes a reduction in both anterograde and retrograde transport (Theisen et al., 2012, Siddiqui et al., 2019). Kinesin-3 motors are thought to be weak motors that work in teams for cargo transport (Okada et al., 1995, Okada et al., 2003, Rogers et al., 2001, Nangaku et al., 1994). Under single molecule conditions, Unc104 and KIF1A, the founding members of the kinesin-3 family, are monomeric and move by biased Brownian motion (Okada and Hirokawa, 1999). This generated force of up to 0.15 pN for each monomeric motor (Okada et al., 2003). However, adding a leucine zipper motif to either truncated or full length Unc104 or KIF1A renders the motor into fast and processive kinesins, which can generate forces of about 3 pN (Budaitis et al., 2021, Tomishige et al., 2002). Because KIF1C is a stable dimer, we hypothesised that single molecules of native KIF1C can generate substantial forces.

Here we determine the force-generating properties of KIF1C and probe the effects of two pathogenic mutations on single molecule motility, force generation and intracellular transport. We find that full-length recombinant KIF1C has a stall force of 5.7 pN and an average detachment force of 3 pN. While HSP-causing mutations P176L and R169W permit fast and processive unloaded motility, force generation is severely reduced with peak forces of 1.3 pN and 0.7 pN respectively. Even though motors can work in teams inside cells, the weakened motors still show significant intracellular transport defects.

## Results

### KIF1C is a strong, processive motor

To determine the force output of KIF1C, we used a classic single bead optical tweezer assay (Block et al., 1990, Carter and Cross, 2005) to probe the force response of full-length recombinant human KIF1C (Siddiqui et al., 2019) and for comparison full-length *Drosophila melanogaster* kinesin heavy chain (KHC) (Coy et al., 1999, Toleikis et al., 2020). In this assay, a single motor protein pulls the bead against the force of the trap until it reaches stall force or detaches. The bead displacement is tracked with nanometer accuracy and forces calculated according to Hooke’s law. As expected, KIF1C takes forwards steps of approximately 8 nm length consistent with a kinesin binding site on each tubulin dimer (Figure S1A-B). Both motors produced forces of similar magnitude and duration, although KIF1C force events were more frequently interrupted by backwards slippage events (Figure 1A). KHC backslips once for every 8.8 forward steps and manages to stop backslips at the next binding site in the vast majority of cases, while Kif1C backslips twice as frequently and three times as many KIF1C backslips are larger than 12 nm, suggesting that KIF1C is both more likely to slip backwards and to slip further when it does so (KHC: 16% of backslips >12nm; KIF1C 37% of backslips >12nm) (Figure 1C). Indeed, we find that the time lag between force generating events follows a triexponential distribution, which presumably reflect long backslips to the trap centre for the fastest timescale (31% of KIF1C events with t½ = 74 ms; 13% of KHC events with t½ = 104 ms), and detachments with two different probabilities of reattachment and commencing another productive force event (Figure S1C). Considering long backslips to the trap centre, KIF1C backslips on average at every second step and KHC once every seven steps. As the ratio of detachments to forward steps is also higher in KIF1C, the average duration of KIF1C force events is reduced by 35% compared to KHC (KHC 1.23 ± 0.09 and KIF1C 0.80 ± 0.02, mean ± SEM) (Figure S1D). Under the conditions of our assay, the stall force of KHC, i.e. the force at which the probability of taking a forwards step or slipping backwards is equal, was 7.3 ± 0.2 pN (mean ± SD), while the average detachment force was 4.1 ± 0.07 (mean ± SEM) pN (Figure 1D-E). These values are in agreement with published literature (Khalil et al., 2008, Carter and Cross, 2005, Budaitis et al., 2021, Svoboda et al., 1993, Yildiz et al., 2008, Ray et al., 1993). KIF1C generated slightly lower force and stalled at 5.7 ± 0.2 pN (mean ± SD), and its average detachment force was 3.0 ± 0.04 pN (mean ± SEM) (Figure 1D-E). For comparison, the detachment forces of constitutively active truncated KIF1A and Unc104 were reported as 2.7 and 2.4 pN, respectively (Budaitis et al., 2021), while native KIF1A is only weakly processive with a stall force of 0.15 pN (Okada et al., 2003). Free-running beads with a single KIF1C move at about 1 µm/s, but its average forward speed decreases under the load of the trap and reaches zero at 7pN. In comparison, KHC can maintain an average speed of 450 nm/s under force of up to 2.5 pN before slowing down rapidly and forward speed is zero at its stall force (Figure 1F, Figure S1E).

**Figure 1:**
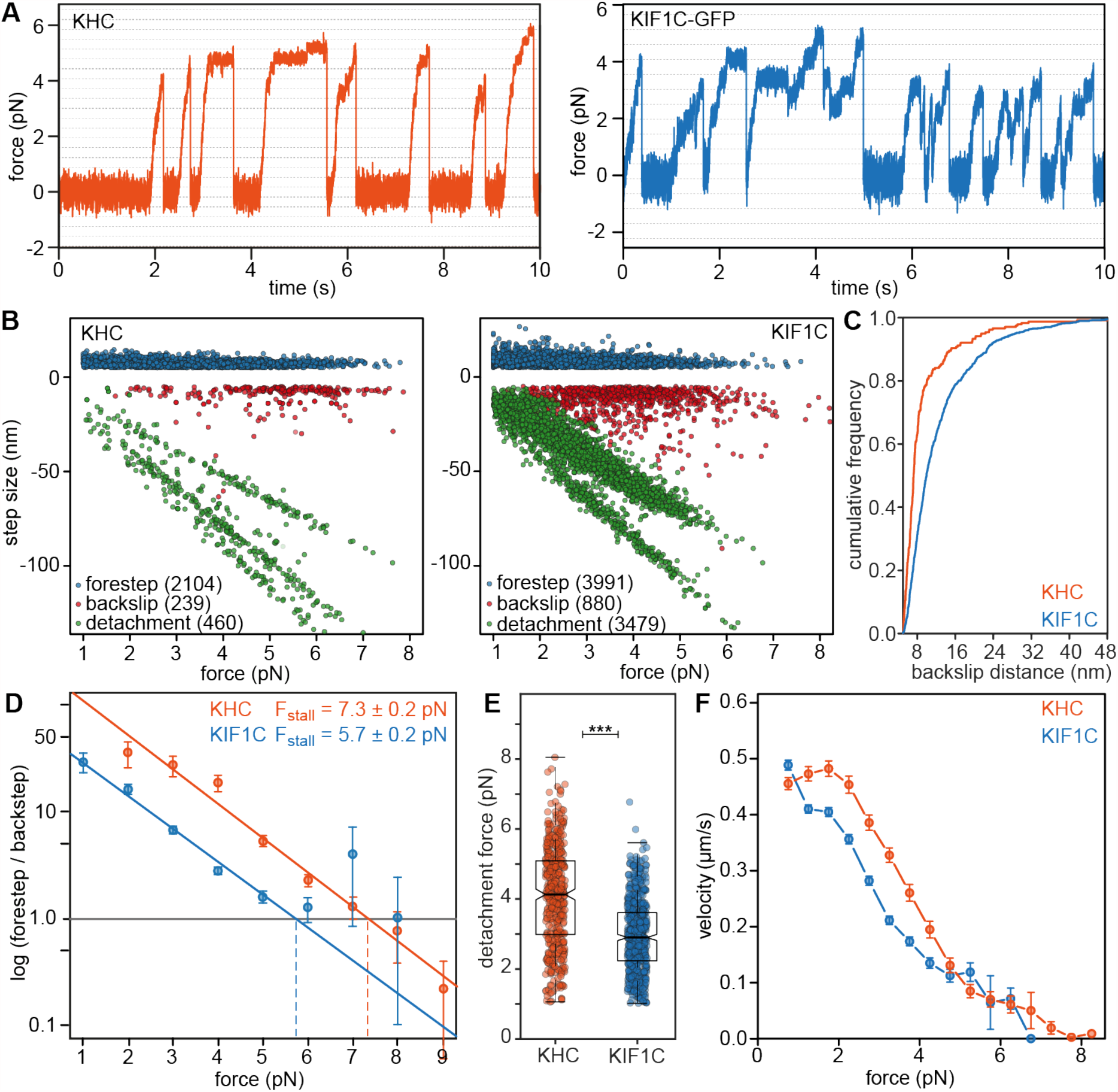
KIF1C single molecule force generation. **(A)** Representative traces from single bead optical trapping experiment for Drosophila melanogaster kinesin heavy chain (KHC) and human KIF1C. Dotted lines indicate 8 nm displacement. **(B)** Step sizes extracted from single molecule force recordings of KHC and KIF1C plotted relative to the force at which they originated. Number of events for each category are given in brackets. **(C)** Cumulative distribution of backslip distances. n=239 for KHC and 880 for KIF1C. p=2•10^−21^ (Kolmogorov-Smirnov test). **(D)** Forestep to backslip ratio relative to force shows 5.7 ± 0.2 pN stall force for KIF1C and 7.3 ± 0.2 pN stall force for KHC. Note that superstall events obtained under added resistive loads are only included in the KHC dataset. **(E)** Distribution of detachment forces. n=460 for KHC and 599 for KIF1C. *** p<0.0001 (t-test). **(F)** Force-velocity relationship for KHC (n=353 individual runs) and KIF1C (n=1662).

Thus KIF1C is not only a highly processive motor (Siddiqui et al., 2019), but also generates substantial forces, which might explain why KIF1C translocates cargoes in cells more efficiently than any other kinesin (Lipka et al., 2016).

### Pathogenic mutations increase off rate and speed of KIF1C

In recent years, disease-causing mutations have been identified for a number of patients representing with complicated forms of hereditary spastic paraplegia or spastic ataxias. Two of these, P176L (Caballero Oteyza et al., 2014) and R169W (Dor et al., 2014) are located in the motor domain of KIF1C close to the microtubule binding interface (Figure 2A), but how these affect the biophysical properties of KIF1C is currently unclear. Thus, we introduced both mutations and purified full-length recombinant human KIF1C-GFP with each mutation from insect cells (Figure 2B). To investigate the effect of these pathogenic mutations, we first determined in single molecule assays whether the motors can still bind and move along microtubules. Both KIF1C patient mutant proteins, KIF1C_P176L_-GFP and KIF1C_R169W_-GFP were motile in the single molecule motility assays and showed microtubule plus-end accumulation similarly to the wildtype (Figure 2C), suggesting that the mutants are still able to move processively along microtubules. Under the conditions of the assay, we observe slightly more than half of motors being static or moving very slowly (< 25 nm/s). The fraction of static motors was slightly increased for both mutants (wt: 53 ± 5 %; P176L: 65 ± 9 %; R169W: 61 ± 7 % (mean ± SD)). Diffusing motors are observed occasionally for wildtype KIF1C, but were increased more than two-fold for the mutants (wt: 3.9 ± 3.9 %; P176L: 8.2 ± 3.4 %; R169W: 8.6 ± 4.5 % (mean ± SD)). Taken together, this resulted in a ∼30% reduction in motile motors (wt: 43 ± 6 %; P176L: 27 ± 6 %; R169W: 30 ± 10 % (mean ± SD)) (Figure 2D-E). Detailed analysis of single molecule motility revealed that the dwell time of both mutants was significantly reduced (wt: 36.3 ± 1.4 s; P176L: 25.8 ± 1.6 s; R169W: 16.0 ± 1.3 s (mean ± SEM)) (Figure 2F). As both mutations are located at the microtubule binding interface of KIF1C, the increase in diffusive events and faster off rates suggest that microtubule binding of both mutants is reduced. Even though the speed was increased (wt: 284 ± 15 nm/s; P176L: 319 ± 29 nm/s; R169W: 362 ± 24 nm/s (mean ± SEM)) (Figure 2G), the fast off rate resulted in significantly shorter run length of both mutants compared to wild type KIF1C (wt: 5.08 ± 0.19 µm; P176L: 3.44 ± 0.15 µm; R169W: 2.80 ± 0.18 µm (mean ± SEM)) (Figure 2H). KIF1C motors frequently pause during runs (Figure 2D), the duration of run phases was more strongly reduced (−24% for P176L and -58% for R169W compared to wt) as a result of the mutations than the duration of pauses (not significantly changed for P176L and -32% for R169W compared to wt) (Figure S2), suggesting that motors are more prone to detachment during runs. Thus, our single molecule analysis of full length KIF1C wildtype and P176L and R169W mutants suggests that both pathogenic mutations reduce microtubule binding affinity, which results in faster motility, but a higher off rate and shorter run length.

**Figure 2:**
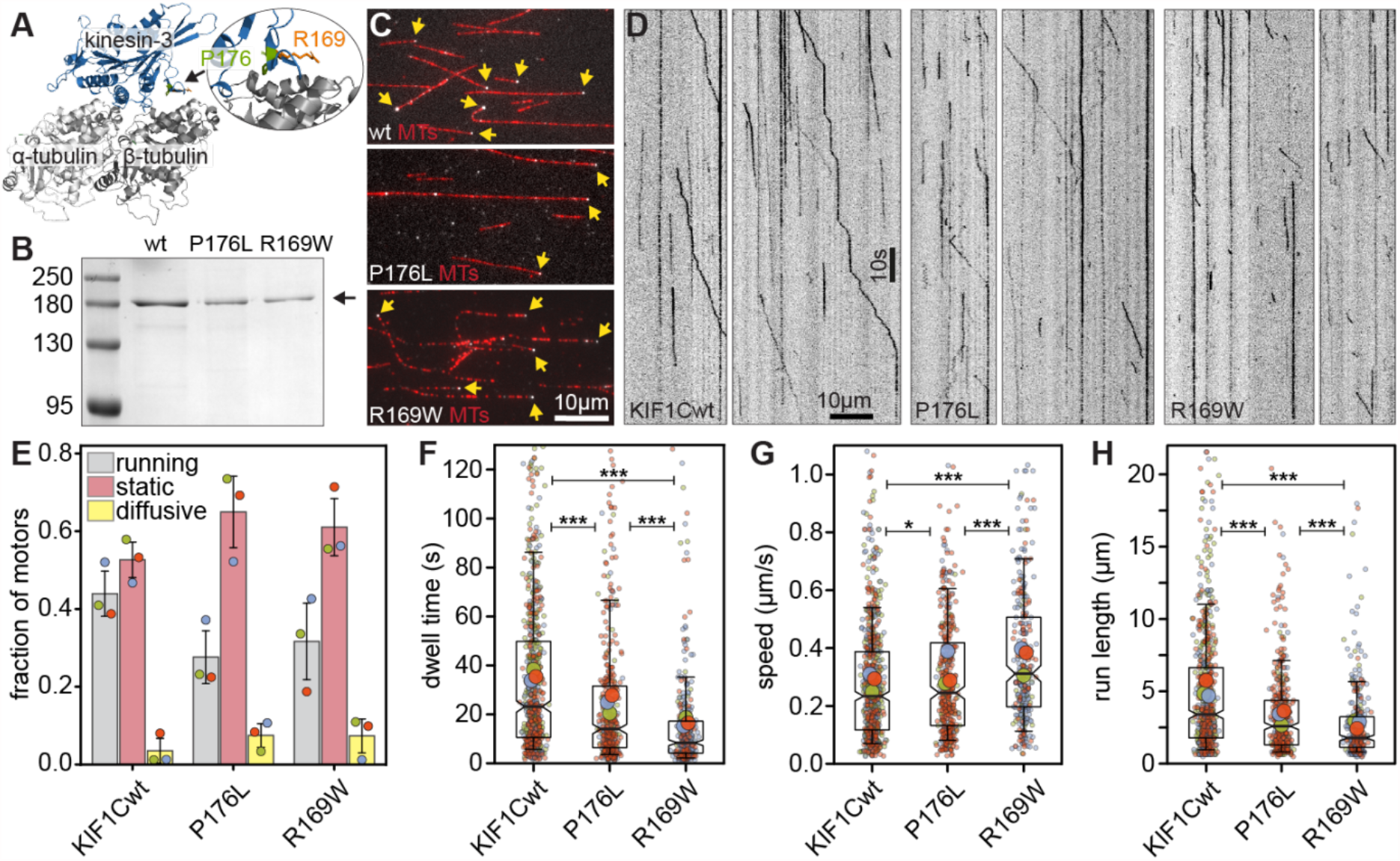
Pathogenic mutations in KIF1C increase unbinding rate and speed. **(A)** Structure of a kinesin-3 motor domain (blue) bound to a microtubule (grey) with the residues R169 and P176 highlighted. Structure is of related motor KIF1A (PDB reference 4UXP) in which both residues are conserved. **(B)** SDS PAGE analysis of purified recombinant human KIF1C-GFP wildtype, KIF1C_P176L_-GFP and KIF1C_R169W_-GFP and molecular size markers as indicated. **(C)** TIRF micrograph of Taxol-stabilised microtubules (red) and full-length recombinant human KIF1C tagged with GFP (wildtype or carrying mutations as indicated) in greyscale. Plus end accumulation is indicated with yellow arrows. **(D)** Representative kymographs showing motility of KIF1C-GFP wildtype and pathogenic mutants P176L and R169W. **(E)** Fraction of motors that run towards the plus end, are static or diffusive. Bars show mean ± SD of 3 experiments. Coloured dots indicate individual values for each experiment. **(F-H)** Superplots for dwell time, speed and run length of individual motors (small dots), averages per experiment day (large dots) are shown colour-coded by experiment day. Boxes indicate quartiles and whiskers at 10/90%. Note that values outside of the Y axis limits have been omitted, but are included in the statistics. * p<0.05; *** p<0.0001 (Kruskal Wallis, Conover’s test post hoc with Holm correction).

### Pathogenic KIF1C mutants are impaired in force generation

In cells, motors haul cargoes against the viscous drag of the cytoplasm and various intracellular structures. Thus, to understand how the reduced microtubule affinity affects the ability to step under load, we analysed both KIF1C_P176L_-GFP and KIF1C_R169W_-GFP using the optical trap. At the trap stiffness we used to analyse KIF1C wild type motors (Figure 1), we did not observe significant bead displacements for the mutants. To confirm that motors were present on the beads, we positioned beads on the microtubule and then switched the trap off to let the motor run without load. This confirmed the presence of functional motors (Figure 3A), but these were too weak to generate forces comparable to the wildtype motor. We then collected bead displacement data at reduced trap stiffness (see methods for details) and detected short runs for KIF1C_P176L_-GFP with an average peak force of 1.3 ± 0.03 pN (mean ± SEM), compared to 3.5 ± 0.03 pN (mean ± SEM) for the wildtype motor (Figure 3B-C). KIF1C_R169W_-GFP moved the bead only small distances when the trap was switched off and could not generate forces above 1 pN (Figure 3B-C). These data suggest that the moderate reduction in processivity of unloaded motors observed in single molecule assays are exacerbated once the motors have to generate forces against a load.

**Figure 3:**
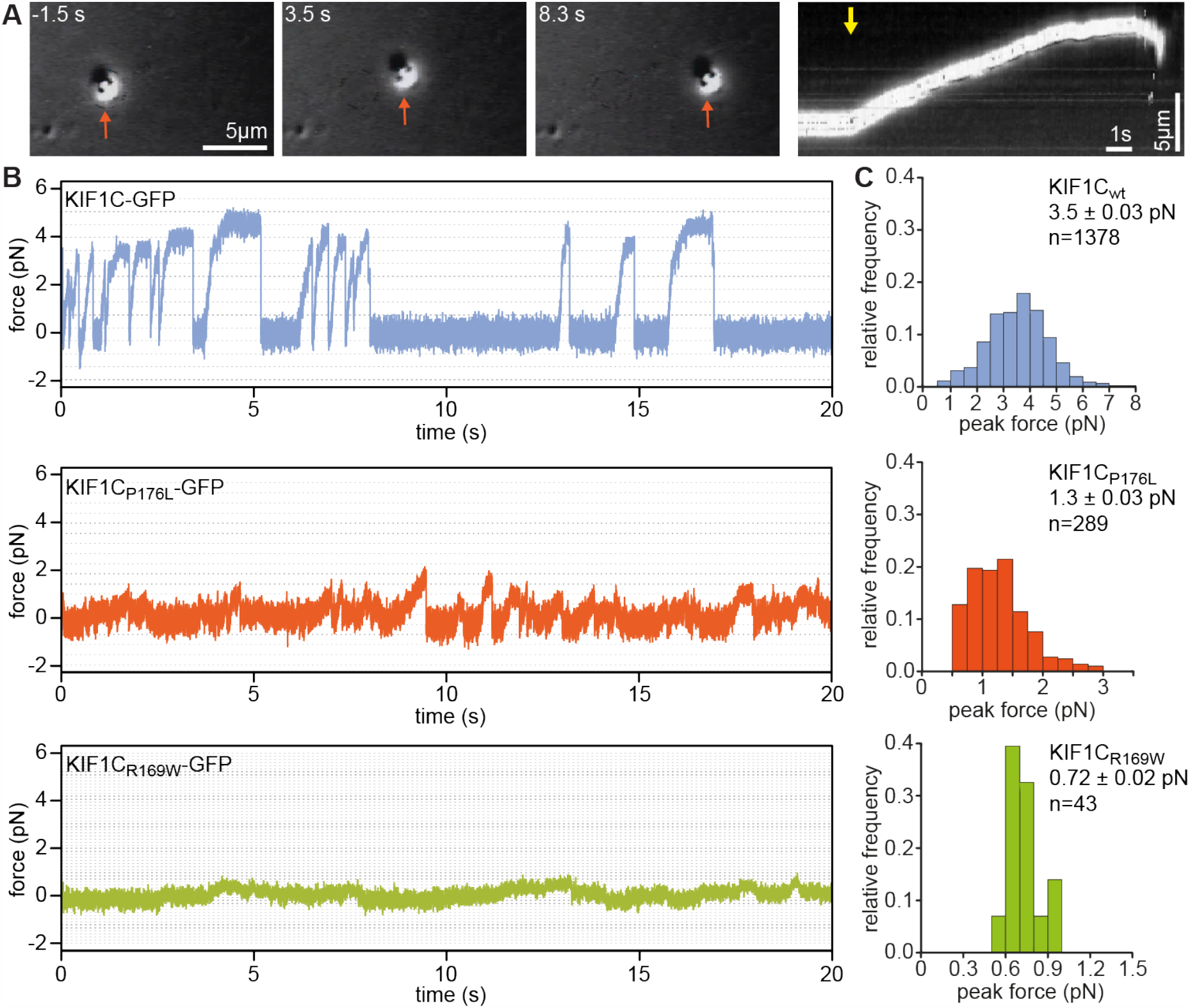
Pathogenic mutations of KIF1C impair force generation. **(A)** Stills and kymograph from an optical trapping experiment showing processive bead transport by KIF1C_P176L_-GFP after release of the motor from the trap (bead indicated with orange arrow in stills, indicated time relative to trap release indicated by yellow arrow in kymograph). **(B)** Representative traces from optical trapping experiment with KIF1C-GFP wildtype and pathogenic mutants P176L and R169W. Dashed lines indicate 8 nm displacement intervals, which are narrower at low trap stiffness used for mutants. **(C)** Histograms of peak forces determined for event lasting longer than 200 ms that exceed 0.5 pN. Mean ± SEM and number of events are indicated. Groups are significantly different with p<0.0001 (Kruskal Wallis, with Sidak correction).

### Pathogenic mutations impair cellular cargo transport

In cells, motors could work in teams and thereby might be able to compensate for the reduction in force generation on the single molecule level. In order to test this, we first analysed the ability of KIF1C with the pathogenic mutations to localise to the cell periphery, which would indicate that they are able to move through the cytoplasm, presumably while carrying cargoes.

KIF1C-GFP accumulates in the tails of migrating RPE1 cells as previously reported (Theisen et al., 2012, Siddiqui et al., 2019). As the extent of the tail accumulation depends on whether the tail is currently growing, stable or retracting (Theisen et al., 2012), we used KIF1C-mCherry as a marker for tail status and co-expressed KIF1C-GFP, KIF1C_P176L_-GFP and KIF1C_R169W_-GFP in RPE1 cells to test their ability to localise to the cell tail. Even though endogenous and wildtype KIF1C-mCherry were present and might allow for heterodimerisation with the mutants, we found that tail accumulation of KIF1C_R169W_-GFP (1.02 ± 0.03 (mean ± SEM)); was significantly reduced compared to wildtype KIF1C-GFP (1.43 ± 0.04 (mean ± SEM)). Surprisingly, KIF1C_P176L_-GFP accumulated stronger in tail tips (1.66 ± 0.05 (mean ± SEM)) than the wildtype (Figure 4A-B). One possible explanation could be that the P176L mutation weakens the autoinhibitory interaction between stalk and motor domain as a stalk deletion mutant also over-accumulated in cell tips (Siddiqui et al., 2019). However, we didn’t detect a significant increase in KIF1C_P176L_-GFP landing rate in our *in vitro* experiments, which is incnsistent with the idea of hyperactivation. An alternative explanation is that KIF1C_P176L_-GFP is not arriving at a higher rate at cell tips, but is retained more strongly or fails to efficiently participate in retrograde transport.

**Figure 4:**
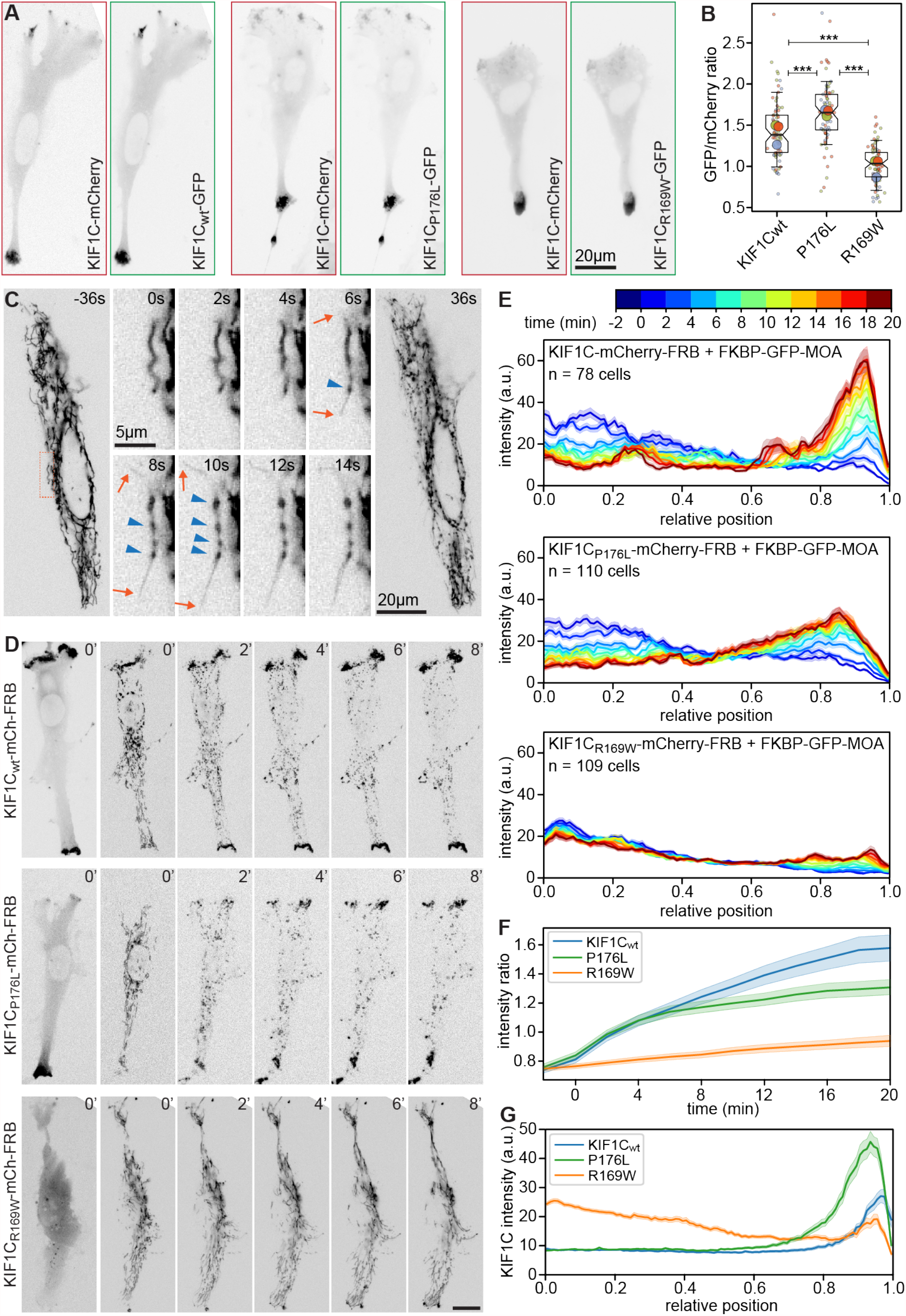
Pathogenic mutations of KIF1C impair intracellular cargo transport. **(A)** RPE1 cells co-expressing KIF1C-mCherry and either wildtype KIF1C-GFP, KIF1C_P176L_-GFP or KIF1C_R169W_-GFP. All proteins accumulate in the tip of cell tails if overexpressed. **(B)** Superplot of 3 cell tail accumulation experiments showing ratio of enrichment in cell tails versus perinuclear cytoplasm for GFP and mCherry constructs. Small circles represent one cell and large circles mean for experiment. Box plots show quartiles and 10/90% whiskers of pooled data. *** p<0.0001 (Kruskal Wallis, Conover’s test post hoc with Holm correction). **(C)** Mitochondria morphology changes upon KIF1C recruitment in RPE1 cell expressing KIF1C-mCherry-FRB and FKBP-GFP-MOA. GFP channel shown, indicated times relative to rapamycin addition. Orange box indicates location of zoomed region. Orange arrows indicate membrane tubules pulled out, blue arrowheads indicate beads formed. **(D)** RPE1 cell co-expressing FKBP-GFP-MOA and either wildtype KIF1C-mCherry-FRB, KIF1C_P176L_-mCherry-FRB or KIF1C_R169W_-mCherry-FRB. Left panel shows KIF1C signal, other panels MOA-labelled mitochondria. Time from rapamycin addition indicated in minutes. Scale bar 20µm. **(E)** Averaged linescans showing mitochondria redistribution upon recruitment of wildtype KIF1C, P176L or R169W mutant. Cell position normalised to 0 in cell centre and 1 at cell periphery. Errors show SEM. Time colour-coded as indicated. **(F)** Ratio of GFP-MOA intensity at the cell periphery versus perinuclear area over time. Errors show SEM. **(G)** Averaged linescans showing KIF1C recruitment and KIF1C distribution before rapamycin addition in the same cells analysed for mitochondria redistribution. Cell position normalised to 0 in cell centre and 1 at cell periphery. Errors show SEM.

To more directly test force generation in cells, we recruited wildtype and mutated motors to mitochondria as a relatively large organelle that would be expected to generate a significant amount of viscous drag in the cytoplasm, but also allow large teams of motors to work together. To do this, we co-expressed FKBP-GFP-myc-MAO with either KIF1C-mCherry-FRB, KIF1C_P176L_-mCherry-FRB or KIF1C_R169W_-mCherry-FRB. The tagged monoamine oxidase allowed us to visualise normal mitochondria morphology and distribution before recruiting the full-length motors to the outer mitochondria membrane. Recruitment of wildtype KIF1C caused dramatic changes in mitochondria morphology within seconds of adding rapamycin. The pulling forces fragmented mitochondria into thin outer membrane tubules and beads (Figure 4C). Both tubules and beads were transported towards the cell edges efficiently, resulting in a near complete accumulation of mitochondria to the cell tips within the 20 minutes of the experiment (Figure 4D-E). KIF1C_P176L_-mCherry-FRB caused the same mitochondria beading effect albeit more slowly and transported mitochondria to the cell periphery less efficiently than the wildtype motor (Figure 4D-E). KIF1C_R169W_-mCherry-FRB was recruited very efficiently to mitochondria membranes and accumulated on the periphery facing tip of the membrane. Beading was only observed in cells expressing a relatively large amount of KIF1C_R169W_ and then the motor was unable to efficiently move the large (700-1000 nm diameter) cargo. In most cases, normal tubular mitochondria morphology was retained and a subset of these was transported towards the cell periphery (Figure 4D-E). At the beginning of the experiment, the fluorescence signal for the mitochondria marker near the cell periphery was about 25% lower than in the perinuclear area and equal between all three conditions (Figure 4F). Immediately after adding rapamycin, the signal shifted towards the periphery, albeit at a significantly higher rate in cells expressing wildtype KIF1C-mCherry-FRB or KIF1C_P176L_-mCherry-FRB than the R169W mutant. 20 minutes after adding rapamycin, the ratio of peripheral versus centrally located mitochondria was clearly lower compared to wildtype KIF1C (Figure 4F) (wt: 1.58 ± 0.09; P176L: 1.31 ± 0.05, p=0.01; R169W: 0.94 ± 0.04, p=3•10^−11^ (mean ± SEM, t-tests with Bonferroni correction)). Analysing the KIF1C distribution at the start of the experiment showed a large cytoplasmic pool and strongly reduced tail accumulation of KIF1C_R169W_-mCherry-FRB and the overaccumulation of KIF1C_P176L_- mCherry-FRB in cells tips (Figure 4G). The discrepancy between KIF1C accumulation in tips versus ability to redistribute mitochondria suggests that even if motors with pathogenic mutations cooperate, they cannot compensate for their reduced force generation abilities.

## Discussion

KIF1C is a highly efficient transporter in cells. Here we describe for the first time its force generating properties. We find that a single KIF1C molecule takes processive ∼8nm steps against the force of the optical trap, stalls at 5.7 pN and detaches at 3 pN on average. Thus KIF1C’s force output is about 25% lower than that of conventional kinesin, but dramatically higher than the < 1 pN detachment forces observed for full length KIF1A and slightly higher than that measured for a constitutively active KIF1A mutant (Budaitis et al., 2021). While we observe a 3.5-fold higher probability of backward slipping for KIF1C, force-generating events are broadly similar to those of conventional kinesin and distinctly different from the 50-fold enhanced slipping observed for constitutively active KIF1A (Budaitis et al., 2021, Lam et al., 2021). We proposed recently that backslips arise when phosphate is released before a forward step is completed, which converts the motor into a weakly bound state that is pulled backwards under the load of the trap. Backwards slipping is stopped when either of the two motor domains in ADP state re-engages at a binding site and releases nucleotide to start another cycle of trying to make a forward step (Toleikis et al., 2020). Kinesin-3s are about tenfold more processive and twice as fast as kinesin-1 (Soppina et al., 2014, Scarabelli et al., 2015). The observations that KIF1C and KIF1A undertake backslips more frequently, suggests that they release phosphate faster, which might be a trade-off for being faster motors. However, due to their interaction with microtubules being significantly stronger (Scarabelli et al., 2015), kinesin-3s stay attached to the microtubule even in a weakly bound state for long enough to wait for the next step. The ability to slip in a weakly bound state might also help kinesin-3 motors to work efficiently in teams as it might permit motors to slip forward. Thus not each motor needs to hydrolyse ATP for every step the team moves and the team can potentially move faster than a single motor (Rogers et al., 2001).

The two pathogenic mutations in the L8 region of KIF1C retain motility of the motor but with reduced run length in line with findings for pathogenic mutations in the microtubule binding interface of KIF1A (Lam et al., 2021, Boyle et al., 2021). However, we find that the unloaded velocity of the motors was not negatively affected and even slightly increased, while mutations in the microtubule binding interface of KIF1A resulted in lower single molecule velocities (Lam et al., 2021, Boyle et al., 2021). This could be due to differences in the mutation sites as S274L, P305L and R316W locate around the kinesin-3 specific K-loop region in α4, 3_10_ helix and α5, while the mutations we studied here are at the β-tubulin facing tip of the motor domain in loop 8. When two L8 mutations (R167S and H171D) were simultaneously introduced into KIF1A, run length was 10-fold and speed two-fold reduced (Scarabelli et al., 2015). Two mutations, E220A and L317A, with slightly higher velocity, but reduced microtubule affinity were identified by alanine scanning mutagenesis in kinesin-1, which did not include the sites studied here (Woehlke et al., 1997). Alternatively, this could be a consequence of the distinct biophysical properties of KIF1A and KIF1C motors. This notion is supported by the observation that KIF1C velocity increases with increasing salt concentration in the assay buffer (data not shown), which presumably reduces microtubule affinity in a similar way as the mutants. Despite retaining processive motility and being able to still reach the plus end of microtubules, the force generation of P176L and R169W mutants was reduced to 1/3 and 1/5 of the wildtype motor, respectively. This demonstrates that these pathogenic KIF1C mutations affect force generation much more dramatically than their unloaded motility. This seems not to be the case for the recently characterised P305L mutation in the 3_10_ helix of KIF1A, which showed a roughly 5-fold reduction in landing rate, a 2-fold reduction in speed and run length and a 2-fold reduction in force generation (Lam et al., 2021). It will require detailed analysis of mutations in different regions of the microtubule binding interface to understand whether these differences are due to a specific function of L8 in limiting the velocity and enhancing force generation of KIF1C or whether the relationship of microtubule affinity and force generation differs between motors from the kinesin-3 family.

Ultimately, it will be important to correlate the biophysical properties of the mutated motors with specific defects in neuronal cargo delivery and organelle distribution, defects in neuronal function and survival, time of onset and the severity of the disease presentation in the patients. This would lead to detailed understanding of the disease aetiology and offer a starting point to develop treatment strategies that delay or indeed reverse the symptoms.

## Materials and methods

### Protein expression and purification

pFastBac-M13-6His-KIF1C-GFP was described previously (Siddiqui et al., 2019) and is available from Addgene (130975). The human patient mutations P176L (Caballero Oteyza et al., 2014) and R169W (Dor et al., 2014) were introduced into this plasmid as follows. *pFastBac-M13-KIF1CGFP*_*P176L*_ was generated using *pFastBac-M13-6His-KIF1C-GFP* as template in a three-step mutagenesis PCR with upstream primer pFB5’ (5’-GATTACGATATCCCAACGACC-3’), downstream primer AS358 (5’-GAAGGGATCCACAGTTCCCCCATCCTC-3’) and mutagenesis primer AS528 (5’-CTGCACGTAC<u>**A**</u>GGCCCAGGATG-3’). The fragment containing the mutation was replaced in *pFastBac-M13-6His-KIF1C-GFP* using *Asc*I and *Stu*I. *pFastBac-M13-6His-KIF1C-GFP*_*R169W*_ was generated similarly using upstream primer pFB5’, downstream primer UT170 (5’-ACTGACCTTCTCCGAGTCC-3’) and mutagenesis primer AS376 (5’-CTGCGGGTC<u>**T**</u>GGGAGCACC-3’). The fragment containing the mutation was replaced in *pFastBac-M13-6His-KIF1C-GFP* using *Asc*I and *Bsi*WI.

Purification of full length human KIF1C in insect cells was performed as described previously (Siddiqui et al., 2019). Briefly, pFastBac-M13-6His-KIF1C-GFP, pFastBac-M13-6His-KIF1C_P176L_-GFP, and pFastBac-M13-6His-KIF1C_R169W_-GFP plasmids were transformed into DH10BacYFP competent cells (Trowitzsch et al., 2010) and plated on LB-Agar supplemented with 30 μg/ml kanamycin (#K4000, Sigma), 7 μg/ml gentamycin (#G1372, Sigma), 10 μg/ml tetracycline (#T3258, Sigma, 40 μg/ml Isopropyl β-D-1-thiogalactopyranoside (IPTG, #MB1008, Melford) and 100 μg/ml X-Gal (#MB1001, Melford). Positive transformants (white colonies) were screened by PCR using M13 forward and reverse primers for the integration into the viral genome. The bacmid DNA was isolated from the positive transformants by the alkaline lysis method and transfected into SF9 cells (Invitrogen) with Escort IV (#L-3287, Sigma) according to the manufacturer’s protocol. After 5–7 days, the virus (passage 1, P1) was harvested by centrifugation at 300 × g for 5 min in a swing out 5804 S-4-72 rotor (Eppendorf). Baculovirus infected insect cell (BIIC) stocks were made by infecting SF9 cells with P1 virus and freezing cells before lysis (typically around 36 h) in a 1° cooling rate rack (#NU200 Nalgene) at -80 °C. P1 virus was propagated to passage 2 (P2) by infecting 50 ml of SF9 (VWR, #EM71104-3) culture and harvesting after 5–7 days as described above. For large-scale expression, 500 ml of SF9 cells at a density of 1–1.5 × 106 cells/ml were infected with one vial of BIIC or P2 virus. Cells were harvested when 90% infection rate was achieved as observed by YFP fluorescence, typically between 48 and 72 h. Cells were pelleted at 252 × g in a SLA-3000 rotor (Thermo Scientific) for 20 min. The pellet was resuspended in 4 ml of SF9 lysis buffer (50 mM Sodium phosphate pH 7.5, 150 mM NaCl, 20 mM Imidazole, 0.1% Tween 20, 1.5 mM MgCl2) per gram of cell pellet, supplemented with 0.1 mM ATP and cOmplete protease inhibitor cocktail (#05056489001, Roche) and lysed using a douncer (#885301, Kontes) with 20 strokes. Lysates were then cleared by centrifugation at 38,000 × g in a SS-34 rotor (Sorvall) for 30 min, or 200,000 × g for 40 min in a T865 rotor (Sorvall). SP Sepharose beads (#17-0729-01, GE Healthcare) were equilibrated with the lysis buffer and the cleared lysate obtained is mixed with the beads and batch bound for 1 h. Next, the beads were loaded onto a 5 ml disposable polypropylene gravity column (#29922, Thermo Scientific) and washed with at least 10 CV SP wash buffer (50 mM sodium phosphate pH 7.5, 150 mM NaCl) and eluted with SP elution buffer (50 mM sodium phosphate pH 7.5, 300 mM NaCl). The peak fractions obtained were pooled and diluted with Ni-NTA lysis buffer (50 mM sodium phosphate pH 7.5, 150 mM NaCl, 20 mM Imidazole, 10% glycerol) and batch bound to Ni-NTA beads (#30230, Qiagen) for 1 h. The beads were loaded onto a gravity column and washed with at least 10 CV of Ni-NTA wash buffer (50 mM sodium phosphate pH 7.5, 150 mM NaCl, 50 mM imidazole and 10% glycerol) and eluted with Ni-NTA elution buffer (50 mM sodium phosphate pH 7.5, 150 mM NaCl, 150 mM Imidazole, 0.1 mM ATP and 10% glycerol). The peak fractions were run on an SDS-PAGE gel for visualisation and protein was aliquoted, flash frozen and stored in liquid nitrogen.

Recombinant Kinesin Heavy Chain from *Drosophila melanogaster* was purified as described previously (Coy et al., 1999). Briefly, BL21 DE3 (Invitrogen) were transformed with plasmid pPK113-6H-DHK (accession # AF053733), and the cells were grown at 37 °C until OD_600nm_ reached 0.5. Expression was induced with 0.4 mM Isopropyl β-D-1-thiogalactopyranoside at 15 °C overnight. Cells were harvested by centrifugation (3000 *× g*, 15 min, RT). Cell pellets were stored at -80 °C. For purification, pellets were thawed on ice, and cells were lysed by sonication in buffer A (50 mM phosphate buffer pH 7.5, 300 mM NaCl, 10% glycerol, 1 mM MgCl_2_, 0.1 mM ATP, and 40 mM Imidazole). The lysate was clarified by centrifugation (100,000 × *g*, T865 rotor, 30 min, 4 °C) and the supernatant was then loaded onto 1 ml HisTrap HP column (Qiagen) at 4 °C. Unbound protein was washed with buffer A containing 90 mM imidazole. Finally, the protein was eluted over 300 mM imidazole gradient (10 column volumes). The peak fractions were run on an SDS-PAGE gel for visualisation and protein was aliquoted, flash frozen and stored in liquid nitrogen.

### Force measurements

560 nm Polystyrene beads (Polysciences) and motor protein were incubated together in 80 mM PIPES pH 7, 2 mM MgSO_4_, 1 mM EGTA, 1 mM DTT, 3 mg/ml D-Glucose, 0.2 mg/ml casein and 1 mM ATP. The concentration of motor was decreased such that only 20-30% of the beads moved. Flow cell coverslips were plasma-cleaned and functionalized with APTES silane. Two coverslips were stuck together with Dow Corning High Vacuum Grease. Two lines of grease were applied to the base coverslip (22 × 50 mm) using a syringe, the top coverslip (22 × 22 mm) was placed, forming a flow cell of approximately 10 μl capacity. For covalent attachment of microtubules, glutaraldehyde (8%) was added to the flow cell and incubated for 30 min. The flow cell was washed with MilliQ water. The microtubules were diluted to the required concentration and introduced to the flow cell and allowed to adsorb onto the surface. The microtubules were incubated for an hour to allow for covalent attachment. Next, 0.2 µl of bead-motor protein solution was diluted in 20 µl of assay buffer composed of BRB80, 1 mM ATP, 0.4 mg/ml casein, 10 µM taxol and 0.4 µl oxygen scavenger mix (1 mg/ml catalase, 5 mg/ml Glucose oxidase, 50% glycerol). The beads were then flown to the cell and microtubules were visualised using in-built differential interference contrast on the Optical Trap setup (Carter and Cross, 2005). The trap was steered to position the bead above the microtubule and the image was projected onto the quadrant photodiode detector. Bead positions were acquired at 20 kHz for 180 s, trap stiffness 0.065 to 0.075 pN nm^-1^ for KHC, 0.06 to 0.067 pN nm^-1^ for wildtype KIF1C-GFP, 0.01 to 0.052 pN nm^-1^ for KIF1C_P176L_-GFP and 0.014 pN nm^-1^ for KIF1C_R169W_-GFP. A calibration for trap stiffness was done each day before commencement of measurements. Measurements were done under conditions when 20% or less beads were running on microtubules. Traces with unusual high force events that suggested more than one motor is engaged were excluded from the analysis. For wildtype KIF1C-GFP, data was pooled from 38 recordings of 7 motile beads from 3 independent experiments. For KIF1C_P176L_-GFP, data pooled from 21 recordings of 6 motile beads from 4 independent experiments. For KIF1C_R169W_-GFP data pooled from 3 recordings of 2 motile beads from 1 experiment. KHC data are a subset of a previously published dataset (Toleikis et al., 2020) with the exception that step analysis only included data acquired at a comparable trap stiffness to KIF1C data and data from superstall conditions were only included for the stall force analysis. Data analysis was carried out using custom code written in R. From the traces generated, a moving window t-test algorithm (Carter and Cross, 2005) was used to identify steps with the following parameters – t-test score threshold=30, minimum step size=5 nm, minimum force=1 pN, moving average, n=20 (1 ms). The size of the window and the threshold are varied to determine accurate steps. When the bead returns to within 20 nm of the centre of the trap, we designated this as a detachment. Other movements towards the trap centre are designated as backslips. The stall force is defined as the force at which the motor undertakes as many forward steps as backslips. We only considered backslips of up to 12 nm for the fore/back ratio as their probability of detection is comparable to forward steps.

In parallel to single-step analysis, we also carried out a broader run-event analysis. This analysis was used for determining the peak force and duration of force-generation events, and the force-velocity curve. Data was smoothed by 2000-point averaging (100 ms) and events above 0.5 pN force, lasting more than 0.2 s, were included. The end of the run was registered when the event returned below 0.5 pN threshold. Force-velocity relationship was determined from linear fits to the displacement versus time data smoothed by 200 points (10 ms) over 0.5 pN force bins.

### Single molecule motility assays

Microtubules were assembled from 5 μl of 18 mg/ml unlabelled pig tubulin, 0.2 μl of 1 mg/ml HiLyte Fluor 647 or X-rhodamine-labelled tubulin (#TL670M, #TL6290M, Cytoskeleton) and 0.5 μl of 0.5 mg/ml biotin tubulin (#T333P, Cytoskeleton) in 15 µl MRB80 (80 mM PIPES pH 6.8, 4 mM MgCl2, 1 mM EGTA, 1 mM DTT) supplemented with 4 mM GTP. The mixture was incubated at 37°C for 90 min before diluting in 80 μl MRB80 supplemented with 20 μM Taxol. Unincorporated tubulin was removed by pelleting microtubules through a glycerol cushion (30% glycerol in MRB80) at 20,238 × g for 12 min at room temperature. The microtubule pellet was resuspended in 80 μl of MRB80 with 20 µM Taxol and stored at 28°C covered from light for use on the same day.

Coverslips (22 × 22) were cleaned by incubating in 2.3 M hydrochloric acid overnight at 60 °C. The next day, coverslips were washed with Millipore water and sonicated at 60 °C for 5 min. The wash cycle was repeated five times. The coverslips were dried using a Spin Clean (Technical video) and plasma cleaned using Henniker plasma clean (HPT-200) for 3 min. Flow chambers were made using clean glass slides (Menzel Gläser Superfrost Plus, Thermo Scientific) and double-sided sticky tape (Scotch 3 M) by placing the cleaned coverslip on the sticky tape creating a 100 μm deep flow chamber. The surface was coated with (0.2 mg/ml) PLL(20)-g[3.5]-PEG(2)/PEG(3.4)-Biotin (50%) (#PLL(20)-g[3.5]-PEG(2)/PEGbi, Susos AG). Biotin-647-microtubules were attached to this surface with streptavidin (0.625 mg/ml) (#S4762 Sigma) and the surface was blocked with κ-casein (1 mg/ml) (#C0406 Sigma).

KIF1C-GFP, KIF1C_P176L_-GFP and KIF1C_R169W_-GFP were diluted in TIRF Assay Buffer (25 mM HEPES-KOH pH 7.2, 5 mM MgSO4, 1 mM EGTA, 1 mM DTT, 10 µM Taxol) supplemented with 0.05% Tween-20, 25 mM KCl and 200ng/ml κ-casein, spun at 100,000 × g for 10 min in an Airfuge (Beckman Coulter) and fluorescence at 507 nm measured using a NanoDrop™ 3300 Fluorospectrometer to confirm comparable concentrations are used in the assay. Proteins were then added to the motility mix (TIRF Assay Buffer supplemented with 5 mM ATP, 5 mM phosphocreatine (#P7936, Sigma), 7 U/ml creatine phosphokinase (#C3755 Sigma), 0.2 mg/ml catalase, 0.4 mg/ml glucose oxidase, 4 mM DTT, 50 mM glucose (#G8270, Sigma), 25 mM KCl, 10 μM taxol, 0.2 mg/ml κ-casein) and flown into the chamber.

Chambers were observed on an Olympus TIRF system using a ×100 NA 1.49 objective, 488 and 640 nm laser lines, an ImageEM emCCD camera (Hamamatsu Photonics) under the control of xCellence software (Olympus), an environmental chamber maintained at 25 °C (Okolab, Ottaviano, Italy). Images were acquired at 5 fps for 180 seconds using 2×2 binning, thus the resulting images have 162 nm pixels and 200 ms temporal resolution.

Motility was analysed by tracing microtubules and generating maximum intensity kymographs from 7 pixel wide lines using the ImageJ Kymograph plugin from Arne Seitz (EPFL Lausanne). Motor tracks from kymographs were traced by hand and recorded as ImageJ ROIs. These paths were analysed in a custom-built python analysis script. Individual phases of tracks were segmented from the ROI, and motors were said to be translocating towards the plus-end or minus-end of the microtubule in each phase if their speed towards that end of the microtubule was greater than 25 nm/s. When the motor’s speed was less than 25 nm/s, it was annotated as paused. Motors with an average speed of less than 25 nm/s or a total run length of less than 500 nm were considered as static. Non-static motors that underwent bidirectional motion of at least 324 nm or just moved towards the minus end where classed as diffusing. Only the remaining, plus end directed motors were analysed for run parameters. Dwell times were calculated as the total time the motor spent on the microtubule until it reaches the plus end or the end of the recording. Run length is the total distance covered by the motor. The average speed was calculated as the run length divided by the dwell time. The pause-corrected average speed was also calculated by dividing the run length by the dwell time minus any time when the motor was paused. Superplots show the mean speed, run-length, or dwell time for a given experimental day, with smaller colour-coded spots showing the individual measured values of motors. On each of the three experiment days, one to four chambers were prepared for each protein. Statistics were calculated at the per-motor level between experimental groups. The data were tested for normality using D’Agostino and Pearson’s test, and as they were not normally distributed, a Kruskal-Wallis test was used to determine if experimental groups differed significantly. Where a statistically significant difference was indicated, pairwise interactions were tested using Conover’s post-hoc test and p-values were corrected for multiple comparisons using the Holm-Bonferroni method.

### KIF1C localisation and mitochondria transport assays

pKIF1C_P176L_-GFP and pKIF1C_R169W_-GFP were based on pKIF1C_RIP2_-GFP (available from Addgene 130977 (Efimova et al., 2014). R169W was introduced using a mutagenesis PCR with upstream primer UT01 (5’-GGAATTC<u>**T**</u>GGAGCTATGGCTGGTG-3’), mutagenesis primer AS376 (5’-CTGCGGGTCTGGGAGCACC-3’) and downstream primer UT170 (5’-ACTGACCTTCTCCGAGTCC-3’). The fragment containing the mutation was replaced in pKIF1C_RIP2_-GFP using *Eco*RI and *Bsi*WI. P176L was introduced using a mutagenesis PCR with upstream primer UT01, mutagenesis primer AS528 (5’-CTGCACGTAC<u>**A**</u>GGCCCAGGATG-3’) and downstream primer AS359 (5’-GGGGATCCCCTCGTTCCCGTTCC-3’). The fragment containing the mutation was replaced in pKIF1C_RIP2_-GFP using *Eco*RI and *Bsp*EI.

pKIF1C-mCherry-FRB, pKIF1C_P176L_-mCherry-FRB and pKIF1C_R169W_-mCherry-FRB plasmids were based on pβactin-KIF1C_MD-GFP-FRB, a kind gift from Caspar Hoogenraad (Lipka et al., 2016) .The KIF1C motor domain was replaced with full length human KIF1C fused to mCherry from pKIF1C-mCherry (available from Addgene 130978 (Theisen et al., 2012)) using *Nhe*I and *Bsr*GI. The mutations were introduced from pKIF1C_P176L_-GFP and pKIF1C_R169W_-GFP using *Nhe*I and *Bam*HI. FKBP-GFP-Myc-MAO was a kind gift from Sean Munro (MRC-LMB, Cambridge) (Wong and Munro, 2014).

hTERT RPE1 cells (ATCC) were grown in DMEM/Nutrient F-12 Ham (#D6421 Sigma) supplemented with 10% FBS (Sigma), 2 mM L-glutamine (Sigma), 100 U/ml penicillin (Sigma), 100 μg/ml streptomycin (Sigma) and 2.3 g/l sodium bicarbonate (#S8761 Sigma) in a humidified incubator at 37°C and 8% CO_2_.

For tail localisation experiments, RPE1 cells were co-transfected with 0.75 µg KIF1C-mCherry and 0.75 µg of either KIF1C-GFP, pKIF1C_P176L_-GFP or pKIF1C_R169W_-GFP using Fugene 6 (Promega) in fibronectin-coated glass-bottom dishes and imaged 24 hours later on an Olympus Deltavision microscope (Applied Precision, LLC) equipped with eGFP, mCherry filter sets and a CoolSNAP HQ2 camera (Roper Scientific) under the control of SoftWorx (Applied Precision). 37°C and 5% CO_2_ were maintained in a TOKAI Hit stage incubator. Images from migrating cells with a tail at the rear expressing moderate levels of both markers were acquired in both channels. To determine tail accumulation, average intensity measurements from the perinuclear area, the tip of cell tails and background outside of the cell were taken for both channels. After background subtraction, the ratio of tail versus cytoplasm intensity was calculated for both channels. The ratio of GFP versus mCherry channel was calculated as the relative enrichment of the protein of interest versus wildtype KIF1C-mCherry. 3 experiments with a total of 69-77 cells were analysed for each condition.

For mitochondria transport experiments, RPE1 cells were co-transfected with 1 µg KIF1C-mCherry-FRB and 0.5 µg FKBP-GFP-Myc-MAO using Fugene 6 (Promega) in fibronectin-coated glass-bottom dishes and imaged 24 hours later on an Olympus Deltavision microscope (Applied Precision, LLC) equipped with eGFP, mCherry filter sets and a CoolSNAP HQ2 camera (Roper Scientific) under the control of SoftWorx (Applied Precision). Live cells were imaged using an Olympus 40x UPlanFL N Oil NA1.3 objective in imaging medium (Leibovitz’s L-15 medium (Gibco™ 21083027) without phenol red, supplemented with 10% FBS (Sigma), 2 mM L-glutamine (Sigma), 100 U/ml penicillin (Sigma), 100 μg/ml streptomycin (Sigma)) at 37 °C. Cells expressing moderate levels of both KIF1C and the mitochondrial marker were identified and their stage position labelled. Images in both channels were acquired from each stage position at 2-minute intervals for 22 minutes to determine organelle redistribution. After the first image from each cell, rapamycin was added to the imaging dish at a final concentration of 200 µM. To observe mitochondria morphology change and motility, images from a single cell were acquired at 2 s intervals for 10 minutes and rapamycin added after 30 seconds.

To quantify organelle redistribution, 33 pixel wide line profiles were acquired from the cell centre to the cell tips at each timepoint. The average intensity of the KIF1C signal along the entire line was used to include only cells with comparable expression level across the wildtype and mutant constructs. Positions along line profiles were normalised so that the cell centre was set to 0 and the cell tip to 1. This normalised distance was binned into 100 equal-width bins and intensity values of MAO for each cell were sorted into these bins. The centre of each bin was plotted against the average MAO intensity +/-the standard error of the binned values. To quantify mitochondria redistribution over time, periphery GFP-MOA signal (0.8 to 1.0) was divided by perinuclear GFP-MOA signal (0.0 to 0.2).

## Acknowledgements

The authors gratefully acknowledge Nicholas Carter and CAMDU (Computing and Advanced Microscopy Unit) for access to the optical trap and microscopy support, Alice Bachmann for generating the KIF1C_R169W_-GFP construct and Lewis Mosby for help with curve fitting.

This work was funded by a Research Prize from the Lister Institute of Preventive Medicine and a Wellcome Trust Investigator Award (200870/Z/16/Z) to A.S., a Lister Institute Summer Studentship and a Chancellor’s International PhD Scholarship from the University of Warwick that supported N.S., and a Ph.D. studentship from the MRC Doctoral Training Partnership (MR/N014294/1) that supported A.J.Z. D.R. is supported in part from the University of Warwick. A.T. and R.C were supported by a Wellcome Senior Investigator Award (103895/Z/14/Z).

## Author Contributions

A.S. and N.S. conceived the project. N.S. cloned and purified KIF1C wt and mutants and performed tail localisation experiments. D.R. cloned mammalian expression constructs, performed single molecule experiments & analysed data. A.T. and N.S. performed Kif1C optical trapping experiments and analysed the data. A.T. purified KHC, performed KHC optical trapping experiments and analysed the data. A.S. performed cellular transport assays. A.Z. analysed single molecule motility and cellular transport data. AS wrote the manuscript with contributions from all authors.

## Supplementary Figures

**Figure S1:**
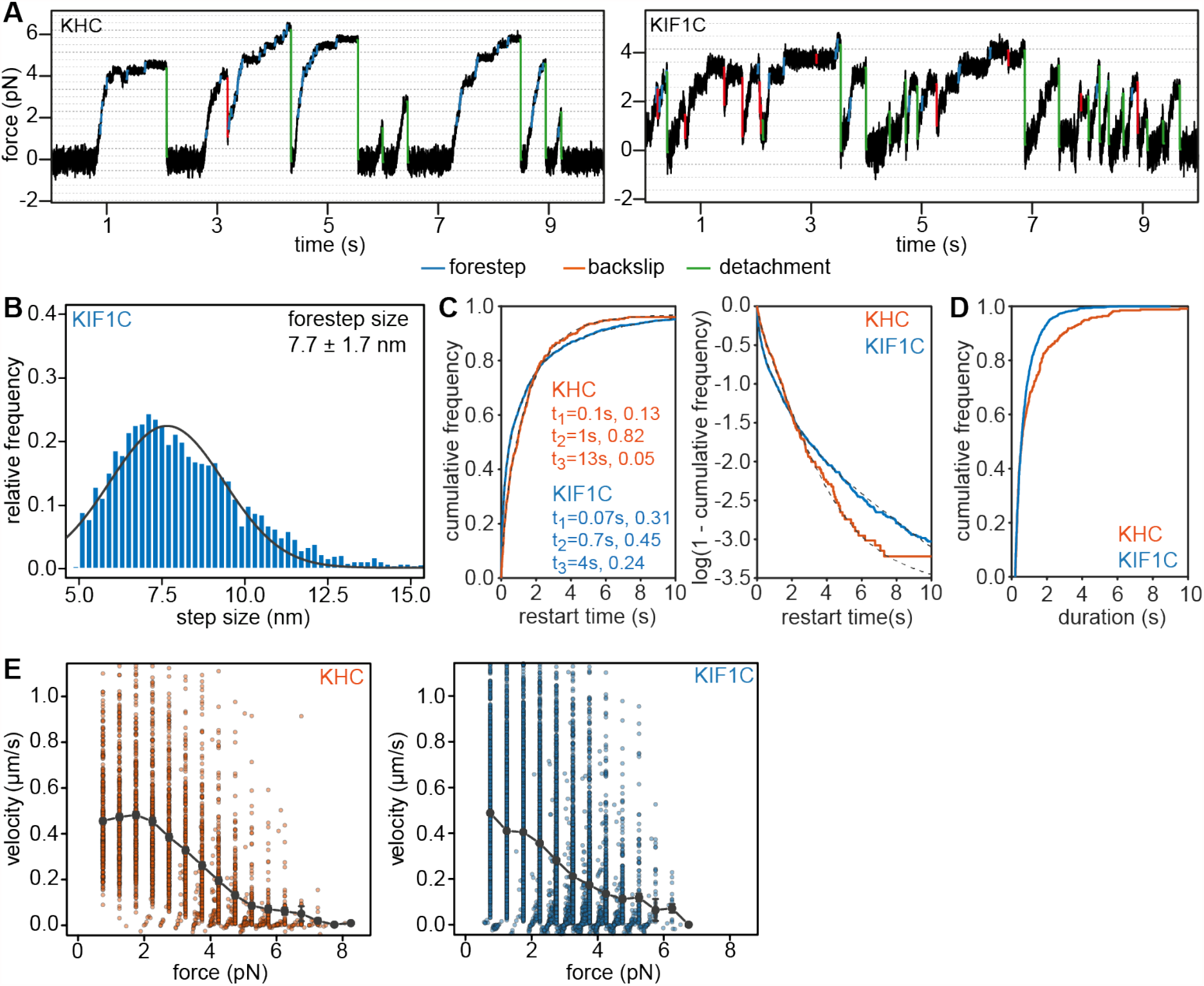
Step and force event analysis. **(A)** Representative traces from single bead optical trapping experiment for Drosophila melanogaster kinesin heavy chain (KHC) and human KIF1C. Dotted lines indicate 8 nm displacements. The steps detected using the step-finder are highlighted. Forward steps in blue, backslips in red and detachments in green. Parameters used in step finder algorithm are t-test score threshold=30, minimum step size=5 nm, minimum force=1 pN, moving average=20 (1 ms). **(B)** Forward step-size distribution for KIF1C. Gaussian curve fit in grey with µ = 7.7 nm, σ= 1.7 nm, n=3991. **(C)** Linear and log plots of cumulative distribution of time lag between force generating events, i.e. time until 0.5 pN force was generated following a backslip or detachment that resulted in force to be reduced below 0.5 pN. n=326 for KHC and n=1404 for KIF1C. Triexponential fits are shown as grey dashed lines and half-times as well as fractions for each mode are indicated in the legend. **(D)** Cumulative distribution of durations over which a force of at least 0.5 pN was maintained. n=333 for KHC and 1378 for KIF1C. * p=0.016 (Wilcoxon rank sum test). **(E)** Distribution of velocity of KHC and KIF1C relative to force. n=353 for KHC and n=1662 for KIF1C.

**Figure S2:**
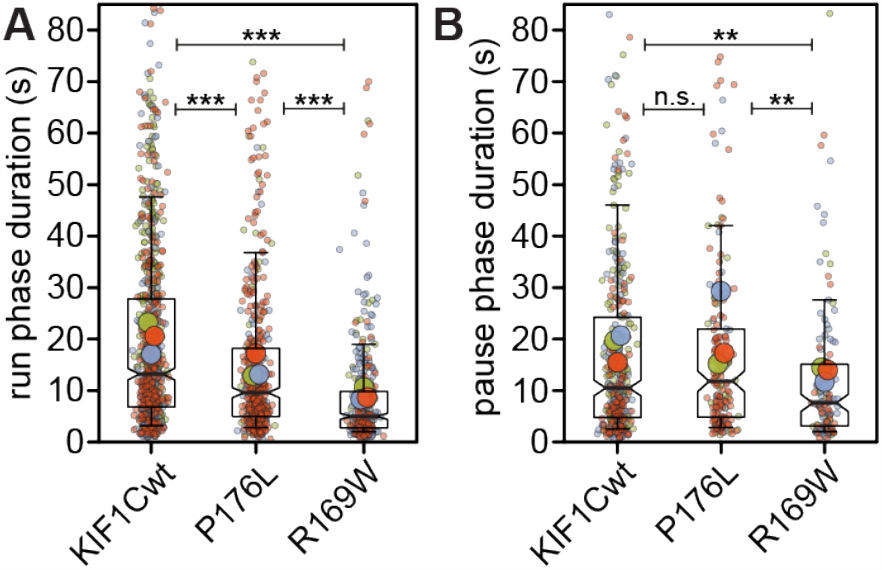
Phase durations from single molecule motility assays. **(A-B)** Superplots for phase durations for plus end-directed runs (A) and pauses (B) of individual motors (small dots), averages per experiment day (large dots) are shown colour-coded by experiment day. Boxes indicate quartiles and whiskers at 10/90%. Note that values outside of the Y axis limits have been omitted, but are included in the statistics. n.s. p>0.5; ** p<0.005; *** p<0.0001 (Kruskal Wallis, Conover’s test post hoc with Holm correction).

